# The antidepressant sertraline provides a novel host directed therapy module for augmenting TB therapy

**DOI:** 10.1101/2020.05.26.115808

**Authors:** Deepthi Shankaran, Anjali Singh, Stanzin Dawa, A Prabhakar, Sheetal Gandotra, Vivek Rao

## Abstract

A prolonged therapy, primarily responsible for development of drug resistance by *Mycobacterium tuberculosis* (Mtb), obligates any new TB regimen to not only reduce treatment duration but also escape pathogen resistance mechanisms. With the aim of harnessing the host response in providing support to existing regimens, we used sertraline (SRT) to stunt the pro-pathogenic type I IFN response of macrophages to infection. While SRT alone could only arrest bacterial growth, it effectively escalated the bactericidal activities of Isoniazid (H) and Rifampicin (R) in macrophages. This strengthening of antibiotic potencies by SRT was more evident in conditions of ineffective control by these frontline TB drug, against tolerant strains or dormant Mtb. SRT, could significantly combine with standard TB drugs to enhance early pathogen clearance from tissues of mice infected with either drug sensitive/ tolerant strains of Mtb. Further, we demonstrate an enhanced protection in acute TB infection of the highly susceptible C3HeB/FeJ mice with the combination therapy signifying the use of SRT as a potent adjunct to standard TB therapeutic regimens against bacterial populations of diverse physiology. This study advocates a novel host directed adjunct therapy regimen for TB with a clinically approved anti-depressant to achieve quicker and greater control of infection.

## Introduction

The current TB therapy regimen ranging between 6 months for pulmonary and 1-2 years for extra pulmonary infections, is often associated with severe drug induced toxicity in patients. Moreover, its failure to completely eradicate the pathogen from the host, forms an ideal platform for the emergence of drug resistant strains (1, 2). It is not surprising that these strains have emerged at an alarming rate in the population and are imposing serious impediments to TB control programs globally (3, 4). Introduction of newer modalities like host directed therapy (HDT) with the potential to reduce duration of therapy and not be affected by pathogen resistance mechanisms offer significant advantages in this scenario (5, 6). Effective molecular entities like antibodies, cytokines, cell based therapies, repurposed drugs have been tested against bacterial and viral infections (7–15). Several facets of infection response ranging from enhancing pathogen clearance to augmenting host metabolism or nutrition have been tapped to develop novel host targeted interventions strategies against complex bacterial infections (16–21).

Mtb infection invokes several mechanisms of pathogen clearance in host cells involving, the induction of pro-inflammatory response, metabolic stress, phago-lysosomal lysis programs, apoptosis/ autophagic mechanisms (22–24). On the other hand, by virtue of its long standing association with humans, Mtb has evolved complex and intricate mechanisms to survive and establish optimal infection in the host (25–32). While attempts to boost the host immune mechanisms for better control of the pathogen are promising, efforts have focused on the development of counter-measures against pathogen mediated subversion of cellular clearance mechanisms (33–38).

We hypothesized that neutralizing a prominent pathogen-beneficial response would indirectly supplement host immunity facilitating better pathogen control. The early, robust, type I IFN response of phagocytes to intracellular bacterial infections including Mtb, is often associated with a detrimental effect on host immune activation and survival (39–42). We sought to offset this response by using sertraline (SRT) - a previously identified antagonist of poly I:C mediated type I IFN signaling, in macrophages and evaluate Mtb infection dynamics (43). We demonstrate that SRT, effectively inhibited infection induced IFN that manifested as growth arrest of Mtb in macrophages. Interestingly, SRT could augment mycobacterial killing in the presence of INH (H) and rifampicin (R), two of the frontline TB drugs in macrophages by effectively lowering the concentration of antibiotics required to achieve clearance. Remarkably, the combination proved effective even against dormant bacilli or antibiotic tolerant Mtb strains. Addition of SRT to TB drugs-HR or HRZE (HR+ pyrazinamide, ethambutol) significantly protected infected mice from TB related pathology both by enhancing bacterial clearance and host survival, implying on the usefulness of this combination therapy in both the intensive and continuation phases of anti TB therapy (ATT). Taken together, we report a novel adjunct TB therapy module by repurposing the FDA approved, prescription antidepressant -sertraline.

## Results

Macrophages respond to Mtb infection by elaborating an array of signaling cascades and effector functions with the nucleic acid driven type I IFN response as an active and dominant response to during infection (42, 44–49). We questioned the benefit to Mtb in actively stimulating this response and hypothesized that suppressing this response in cells would alter macrophage infection dynamics. As an initial step in this direction, we chose a FDA approved drug-SRT with reported activity in suppressing type I IFN in cells and observed a dose dependent reduction in this response of macrophages on treatment with SRT (Fig. 1A). While 1μM was minimally inhibitory, 5μM and 10μM of SRT inhibited the response by 35 and 43% respectively. At a dose of 20μM, SRT significantly reduced the response to 1/5th of the untreated values (Fig. 1A). Moreover, while naïve macrophages were permissive for Mtb growth to 10-folds of input by day 5, SRT, when used at this concentration, despite its minimal activity on Mtb *in vitro*, efficiently restricted growth in infected macrophages (Fig. 1B).

**Fig. 1:**
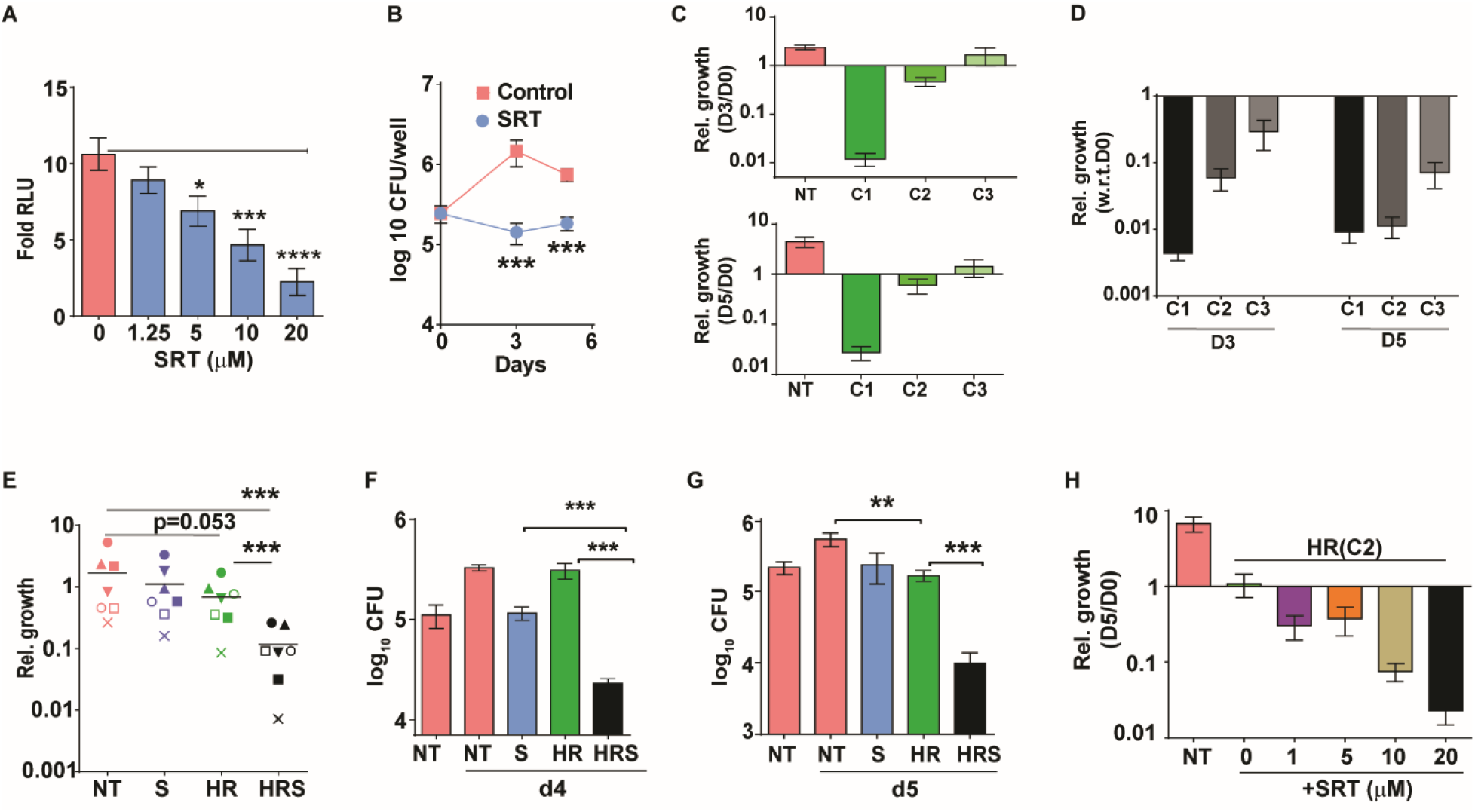
Sertraline inhibits Mtb induced Type I IFN response and restricts intra-macrophage Mtb growth. A) IRF dependent luciferase activity in THP1 Dual macrophages following infection with Mtb at a MOI of 5. Cells were left untreated or treated with increasing concentrations of SRT for 24h in culture and the luminescence in culture supernatants was measured and is represented as mean ± SEM from 2 independent experiments with triplicate wells each. B-H) Intracellular bacterial numbers in THP1 Dual macrophages following infection with Mtb at MOI5 for 6h and then either left untreated (NT-red) or treated with, Sertraline (SRT/ S-blue), (HR-200ng/ml INH and 1000ng/ml Rif-green) or a combination of all three (HRS-black). B-comparison of bacterial growth in untreated or SRT treated macrophages, C) growth of Mtb in cells not treated or treated with HR at 200ng/ml INH and 1000ng/ml Rif (C1), ten-fold lower (C2) and 25-fold lower (C3) concentrations. The relative bacterial counts CFUs at day 3 and day 5 post infection with respect to day 0 (6hp.i.) was calculated and is represented as rel. growth. ± SEM from N=2 or 3 replicate experiments. D) Intracellular bacterial survival in macrophages treated with HR at C1, C2 and C3 with 20μM SRT at days 3 and 5 *p.i*. is represented as growth relative to Day 0+ SEM from N=2 or 3 replicate experiments. E) Mtb growth in primary human M1-differentiated MDMs from PBMC of seven individuals is represented as CFU counts relative to day 0 CFU in untreated samples. Each symbol represents one individual, colours depict the treatment groups as before. F, G) Growth in macrophages treated with Vit. C (F) for 24h post infection or with 200μM Oleic acid for 48h prior to infection (G). H) Macrophages treated with C2 with different concentrations of SRT. Bacterial numbers were enumerated and is represented as average log10 CFU ± SEM from 2-3 independent experiments with triplicate wells each. Except for E, paired t-test comparing ratios, other datasets were compared with unpaired t-test; **p<0.01, ***p<0.001).

This observed bacterial stasis in SRT treated macrophages prompted us to analyze the effect of this treatment in conjunction with frontline antibiotics-INH and Rifamipicin. As an initial step we, analyzed the effectiveness of these two antibiotics in our infection model by using 3 different doses of HR-C1 (200ng/ml INH and 1000ng/ml Rifampicin), C2 (20ng/ml INH and 100 ng/ml Rifampicin) and C3 (8ng/ml INH and 40ng/ml Rifampicin). We enumerated Mtb growth in at day 3 and 5 *p.i.*. As expected, a gradual reduction in antibiotic effectivity was observed with decreasing doses: while C1 was able to control, a 10-fold lower dose of C2 was less effective while the 25-fold lower dose (C3) was ineffective in restricting bacterial numbers from input values (Fig.1C). In contrast to the growth of Mtb in untreated macrophages (2.3x and 4.5x w.r.t. D0), HR treated macrophages at the highest concentration, could effectively control bacterial numbers by ~62 fold by day 3 and ~92 fold by day 5 (Fig. 1C). This ability was reduced to only a 50% reduction (w.r.t. D0) in CFU by the antibiotics at C2. At the concentration of 0.04X (C3), HR failed to control bacterial growth even after 5 days of treatment. Addition of SRT efficiently boosted the bactericidal properties of antibiotics at all concentrations tested (Fig. 1D). Even at the highly effective HR concentration, addition of SRT further reduced bacterial numbers by 2-3 folds. Surprisingly, the ability of SRT to boost antibiotic properties was --- visible at doses C2 and C3. At the C2 dose, SRT significantly lowered numbers by 8 and 50 folds at day 3 and day 5, respectively in comparison to HR. Even at the lowest dose of C3, SRT substantially enhanced bacterial killing by 6- and 20-folds w.r.t HR at days 3 and 5 post infection.

Decreased portioning of potent antibiotics to regions of bacterial presence in the center of granulomas remains a major cause for the improper clearance of bacteria from infected tissues and development of bacterial tolerance and resistance (50, 51). Given the ability of SRT to enhance potencies at ineffective antibiotic concentrations, we evaluated the antibiotic boosting properties of SRT under different physiological conditions that would mirror this *in vivo* situation. As a first step, we verified the ability of SRT to potentiate Mtb control by frontline antibiotics in primary macrophages (human monocyte derived macrophages) from 7 healthy individuals. Although Mtb growth varied across samples, the antibiotic potentiating effect of SRT was preserved (Fig. 1E). Across different individuals, while SRT and HR individually showed minimal but highly variable bacterial control, SRT universally synergized with antibiotics to further reduce bacterial numbers by 10-15 folds supporting a more general adjunct activity of SRT to frontline TB drugs.

Mtb is endowed with the inherent propensity to enter into a drug-tolerant, non-replicating dormant state associated with drug tolerance and antibiotic failure (52). We hypothesized that any advantage in bacterial clearance under such conditions would prove useful in controlling overall infection; warranting the evaluation of SRT in this situation. The recent model of vitamin C treated THP1 macrophages associated with Mtb dormancy and loss of HR efficacy (53) provided an important platform in this direction. Again, while SRT was able to induce bacterial stasis in this model, HR was completely ineffective at controlling bacterial growth. However, HR along with SRT led to 10-12-fold reduction in bacterial numbers within 4 days of treatment indicating its activity against dormant Mtb (Fig. 1F).

More recently, the inability of drugs to distribute equally amongst the spectrum of granulomatous lesions was recognized as an important factor in promoting bacterial tolerance and resistance. The lipid loaded necrotic lesions forming a formidable barrier for entry of frontline TB drugs *in vivo* (54, 55). To mimic lipid rich conditions i*n vitro* studies we treated THP1 macrophages with oleic acid and found poor efficiency of HR in these cells (Fig. 1G). In contrast, a combination of SRT with HR led to more than 10-fold reduction in bacterial loads.

SRT was found to increase HR efficiency in a dose dependent manner. Even at concentrations as low as 1 μM, SRT, could effectively synergize with antibiotics and facilitate growth restriction in macrophages with increased potency of the combination upon increasing the dose of SRT up to 20 μM (Fig. 1H).

Given the ability of SRT to supplement antibiotic properties, we tested if SRT was potent in supplementing the activity of HR in *in vitro* culture. Even upto 6 μM SRT failed to impact Mtb growth either alone or in combination with 3 concentrations of HR in these conditions (Fig. S1). These results hinted at an indirect effect of SRT in enhancing the ability of macrophages to control bacteria rather than a direct benefit to the microbicidal properties of the antibiotics. We used another potent inhibitor of type I IFN signaling – BX795 to understand the effect of restricting this signaling on bacterial growth control by macrophages (Fig. 2 A, B). We first confirmed the ability of BX795 to inhibit IFN in THP1 macrophages in response to Mtb infection and observed complete abrogation even at a very low concentrations of 1.2 μM (Fig. 2A). Treatment with BX795 alone induced a bacteriostasis in Mtb infected macrophages similar to the effect seen with SRT alone (Fig. 2B). However, addition to HR significantly enhanced bacterial killing by 4-5 folds by 3 days of infection again reflecting the properties observed with SRT (Fig. 2C). We reasoned that inhibition of IFN by SRT added the antibiotic properties of frontline TB drugs, then a similar effect of increased antibiotic efficacy would be visible in IFN signaling deficient macrophages. To test this, we compared the intracellular bacterial numbers in cGAS^−/−^ and STING^−/−^ macrophages with WT cells after 3 days of treatment with HR assessed the potentiating effect of negating type I IFN on antibiotic efficacy in the macrophages deficient in IFN signaling (Fig. 2D, 2E). Similar to treatment with SRT, Mtb was restricted in its growth in macrophages with loss of type I IFN signaling as compared to the Wt cells by day 3 of infection. Absence of cGAS enhanced the ability of HR in controlling bacterial numbers by 2-3 folds in comparison to the control observed in IFN sufficient cells (Fig.2D). This effect was amplified in macrophages with a deficiency in STING wherein, the difference in antibiotic mediated control was more than 40-50 folds between the Wt and STING^−/−^ cells (Fig. 2E). A recent study by Teles et.al, 2013 suggested the active suppression of IFNγ dependent-antimycobacterial activity of macrophages as one of the potent pro-pathogenic effects of type I IFN in mycobacterial infections (56). We hypothesized that a reversal of this suppression would lend credence to the role of SRT in reducing type I IFN signaling in macrophages and cells would be more efficient in controlling infection. Indeed, we found 2-fold enhancement in IFNγ mediated control in SRT treated macrophages (Fig.2F).

**Fig 2:**
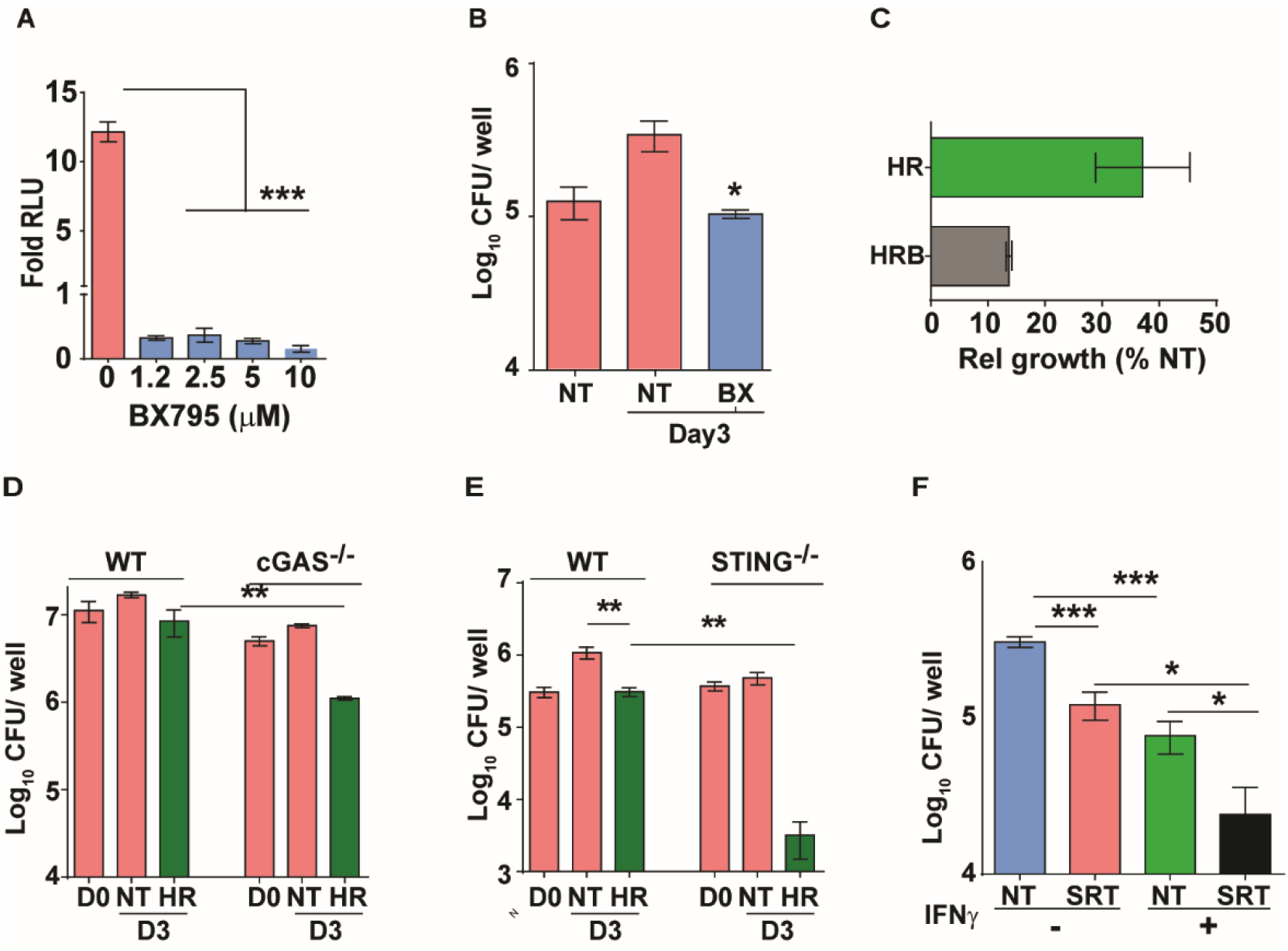
Augmentation property of SRT is due to its ability to inhibit IFN signaling. A) IRF dependent luciferase activity in THP1 dual macrophages 24h after treatment with varying doses of BX795 along with infection with Mtb at MOI of 5. B,C) Bacterial growth in macrophages left untreated or treated with 10μM BX795. Relative growth of Mtb in macrophages treated with HR and HR+BX795 (HRB) for 3 days. The percentage relative growth of intracellular bacterial numbers in HR or HRB groups with respect to untreated samples is depicted. D, E) Bacterial growth in macrophages WT, cGAS^−/−^, STING^−/−^ (G- left untreated or treated with HR), for 3 days. The numbers are mean ± SEM for triplicate wells of N=2 (D) and mean ± SEM for (E) of N=2/3 experiments. F) Growth of Mtb in macrophages pre-treated with IFNγ for 16h, infected with Mtb for 6h and then left untreated or treated with SRT for 3 days. Mean CFU values for triplicate assay wells from two independent experiments (N=2) ± SEM is shown.

The increase in IFNγ mediated bacterial control prompted us to analyze the pro-inflammatory response of Mtb infected macrophages treated with SRT (Fig. 3A-D). Gene expression profiles showed reduction in the expression of TNF and IL1β after 18h of treatment with SRT consistent with reported literature (57). In comparison to HR, while TNF expression was reduced slightly in the HRS treated macrophages (Fig. 3A), negligible levels of TNF was observed in the cell supernatants in SRT treated cells (SRT, HRS), contrasting with the high levels of TNF (between 500pg-1ng/ml) in the case of Mtb infected cells with or without treatment with HR (Fig. 3B). Surprisingly, while expression levels of IL1β were 2-3 folds lower in HRS treated macrophages by 18h (Fig. 3C), the amount of secreted cytokine was significantly elevated (> 2-3 folds) in macrophages treated with SRT/ HRS from 18h until 66h (Fig. 3D). These data strongly indicated potential activation of the host cell inflammasome by SRT. With several studies supporting inverse regulation of type I IFN and inflammasome activation (58, 59), we tested the efficacy of SRT to potentiate antibiotic mediated killing in the presence of inflammasome inhibitors. Again, while SRT enhanced the ability of HR to control Mtb in macrophages, pretreatment of cells with the inflammasome inhibitor, isoliquiritigenin (I), completely nullified the boosting effect of SRT on antibiotic efficacy (Fig. 3E), suggesting that the type I IFN modulating effect of SRT was mediated in part by inflammasome activation.

**Fig. 3:**
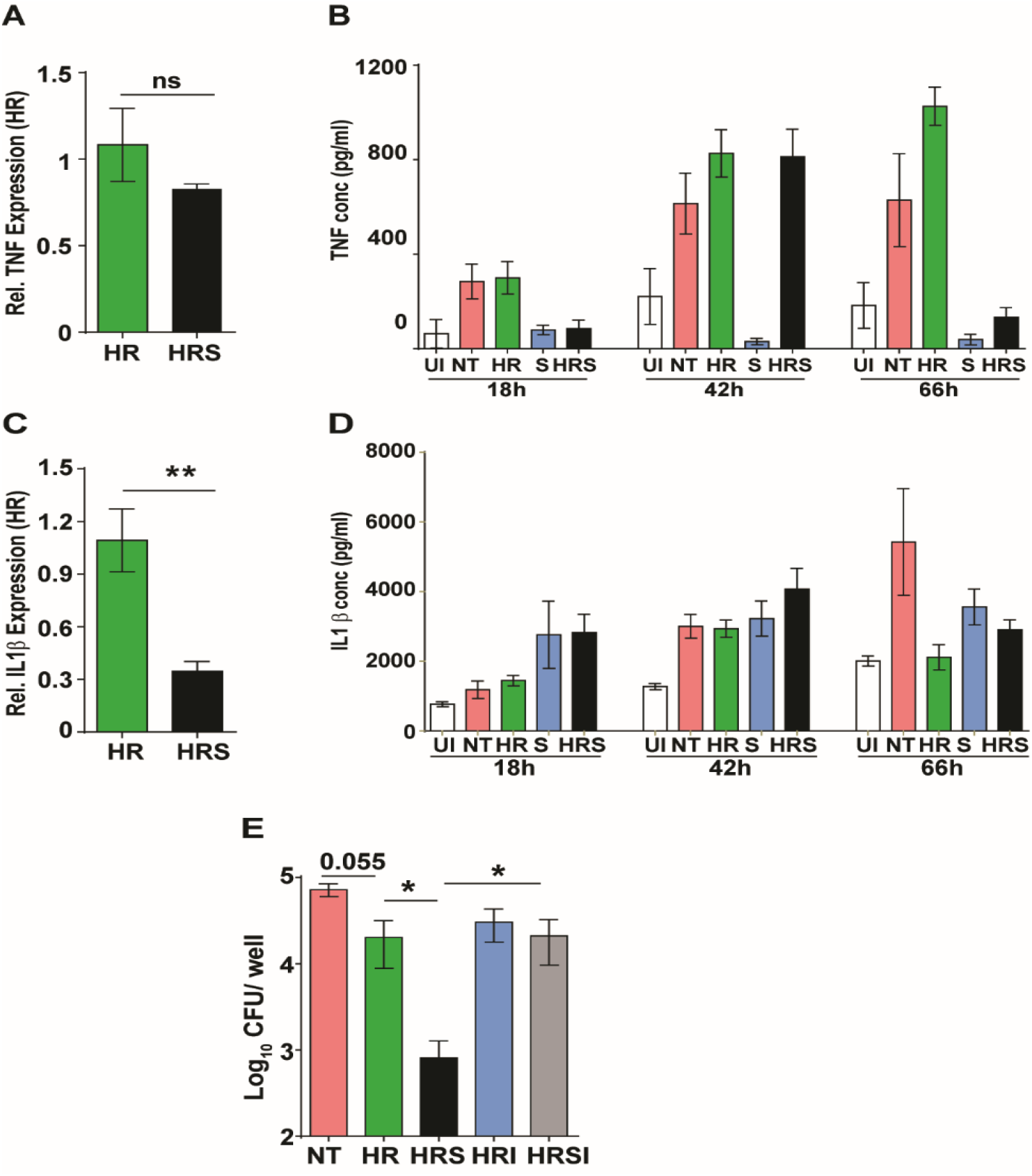
Important role for inflammasome activation in antibiotic potentiation by SRT. Relative expression of inflammatory cytokines in macrophages treated with HR or HRS. Data in A and C depict fold expression (transcript abundance) relative to HR alone as mean ± SEM from two independent experiments with duplicate wells each at 18h post treatment. B and D depict secreted cytokines at indicated time points post treatment. B) Average values ± SEM of TNF at 18h, 43h and 66h post treatment of two independent experiments of triplicate wells (N=2). D) IL1 β levels in cell supernatants at 18h, 42h and 66h of triplicate wells (n=3). E) Growth of Mtb in macrophages infected with Mtb for 6h and then left untreated or treated with HR, SRT, HRS, HRS+ isoliquiritigenin (I) for 5 days. Data is represented as mean ± SEM for 2-3 independent experiments containing triplicate wells per assay and analysed by (unpaired t-test, ns=not significant, **p<0.01, ***p<0.001).

The antibiotic escalating properties of SRT in macrophages prompted testing of this combination regimen *in vivo*. We reasoned that the survival of TB infected C3HeB/FeJ mice would provide an optimal platform for a fast readout of comparative drug efficacies of standard TB drugs and the adjunct regimen with SRT. Given the excellent ability of infection control by INH and Rifampicin in this model, it was necessary to test the adjunct effect of SRT in conditions of minimal advantage imparted by the frontline drug combination alone *i.e.* lower doses -C2 (0.1X) and C3 (0.01X) in addition to the standard dose C1(X) of HR *ad libitum* in drinking water (Fig. 4A). Aerosol delivery of Mtb at a high dose of ~500 cfu/animal, resulted in precipitous disease with rapid killing of untreated (NT) animals by day 31 of infection (Fig. 4B). To facilitate disease progression prior to treatment initiation, animals infected with Mtb were left untreated for 2 weeks. SRT alone was effective in delaying the disease progression in animals as the mean survival time (MST) increased from 31 days for untreated animals to 38 days. HR at the lowest concentration (C3), similar to SRT, deferred animal mortality with MST of 41 days while the higher doses of HR (C2 and C1) increased MST significantly to 85 and >90, respectively. Given the higher susceptibilities of female mice to infection observed in our study, we decided to differentially tally the gender-based effects of SRT treatment in these animals. The combination of HRC3 and SRT, nearly doubled the MST of both female and male mice to 78 days and 100 days from 41 and 45 days respectively (Fig. 4C). This benefit of SRT was similar to that observed for a 10-fold higher concentration of HR alone (MST of 85 for HRC2). The advantage of SRT inclusion with HR was evident in the gross lung pathology by 30 days of infection. Both NT and HRC3 treated lungs showed extensive progressive granulomas, contrasting with a significant amelioration of pathology seen in HR3S treated animals (Fig. 4D).

**Fig. 4:**
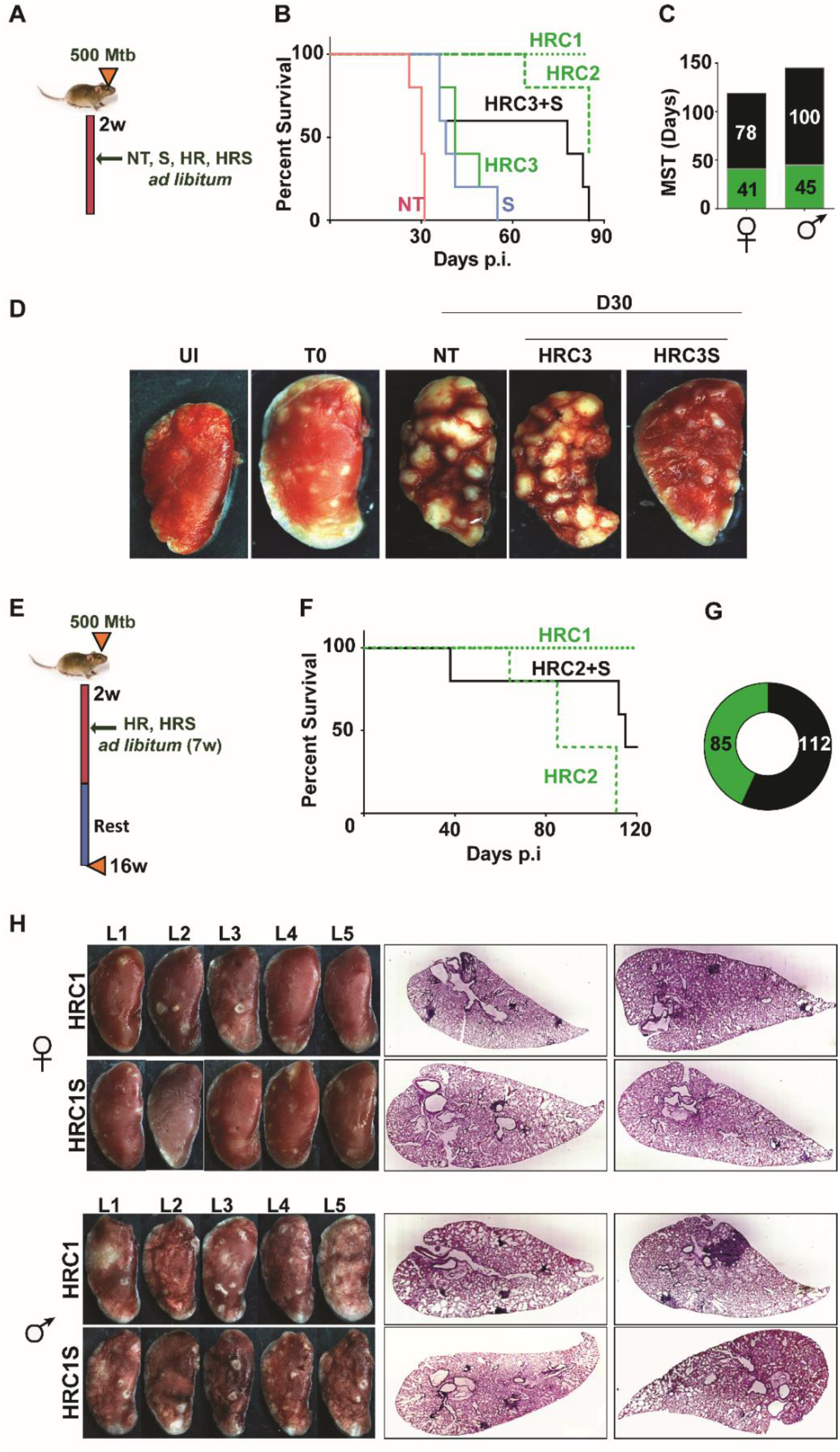
Adjunct SRT improves host survival in a susceptible mouse model of infection. A) Schematic of Mtb infection and drug treatment in C3HeB/FeJ female mice. B) Survival curve of Mtb infected female C3HeB/FeJ mice treated with different concentrations of H and R (HRC1-1X: H-100μg/ml, R-40μg/ml, HRC2-0.1X, HRC3-0.01X) alone or along with SRT (10μg/ml). C) Median survival time of different treatment groups of mice. D) Gross tissue morphology of lungs of uninfected animals (UI) and indicated groups at 30 days post infection with Mtb. E) Schematic of infection and antibiotic treatment in C3Heb/FeJ with HRC2 and HRC1. F-G) Survival (F) and MST (G) of C3Heb/FeJ mice treated with HRC2 or HRC2S. H) Gross lung morphology at the end of 16 weeks and histological sections of lungs with H&E staining of C3HeB/FeJ mice either treated with HRC1 or in combination with SRT.

Further, we wanted to explore temporal benefits of the combination regimen after withdrawal of a limited term treatment (Fig. 4E). In this model, 100% of animals survived after treatment with HR(C1) at 16 weeks post infection (Fig. 4F). A 7-week *ad libitum* treatment with the 10-fold lower dose of HR (HRC2) (Fig. 4F) significantly decreased the MST of animals to 85 days with 100% mortality by 16 weeks. In contrast, 40% of HRC2S treated animals, survived with a MST of 112 days for the group (Fig. 4F, G). All animals with the highest dose of HR (HRC1) either alone or with SRT survived the infection. Despite, the significant heterogeneity of treatment response in male and female mice, with a relatively lower response as evidenced by the greater number of lesions in lungs of male animals, a co-operative effect of SRT inclusion was evident as a significant improvement in TB associated lung pathology (Fig. 4H). Small macroscopic lesions were observed in lungs of 60% of the female mice treated with HRC1 that showed as multiple, well defined granuloma in the H&E stained sections by the 16^th^ week post infection. In contrast, mice treated with SRT and HRC1 showed negligible involvement of the lung tissue in granulomatous cellular accumulation. Even in tissues of male mice, animals receiving the adjunct therapy showed fewer macroscopic and significantly lower numbers of microscopic granuloma in lung sections in comparison to animals treated with the antibiotics alone.

TB treatment in the intensive phase involves the use of 4 frontline TB drugs-HRZE for a period of two months and HR for an additional 4 months. Further to test the efficacy of SRT in combination with HRZE in an acute model of disease, Mtb infected C3HeB/FeJ mice were treated either with the established dose of HRZE or in combination with SRT (Fig. 5A) and evaluated TB associated pathology of the lungs at 16 weeks of infection. Male (Fig. 5B) and female (Fig. 5C) animals treated with the 4 drugs for 7 weeks harbored 10^5^-10^6^ bacteria in their lungs, respectively. Addition of SRT appreciably improved drug efficacy, reducing the lung bacterial loads by a further 5-7 folds (Fig. 5B and C). The drugs efficiently lowered tissue pathology as evidenced by the macroscopic lesions seen in the lungs of infected mice (Fig. 5D). While both female and male mice showed small macroscopic lesions in lungs on treatment with HRZE, despite the heterogeneity between the genders, the combination of SRT and HRZE sufficiently decreased the extent of tissue involvement in TB associated pathology. This difference was more evident in tissue sections, wherein animals treated with HRZES were devoid of granulomatous infiltrates in contrast to the HRZE treated animals which had significantly higher numbers of granulomas in the lungs (Fig. 5D).

**Fig. 5:**
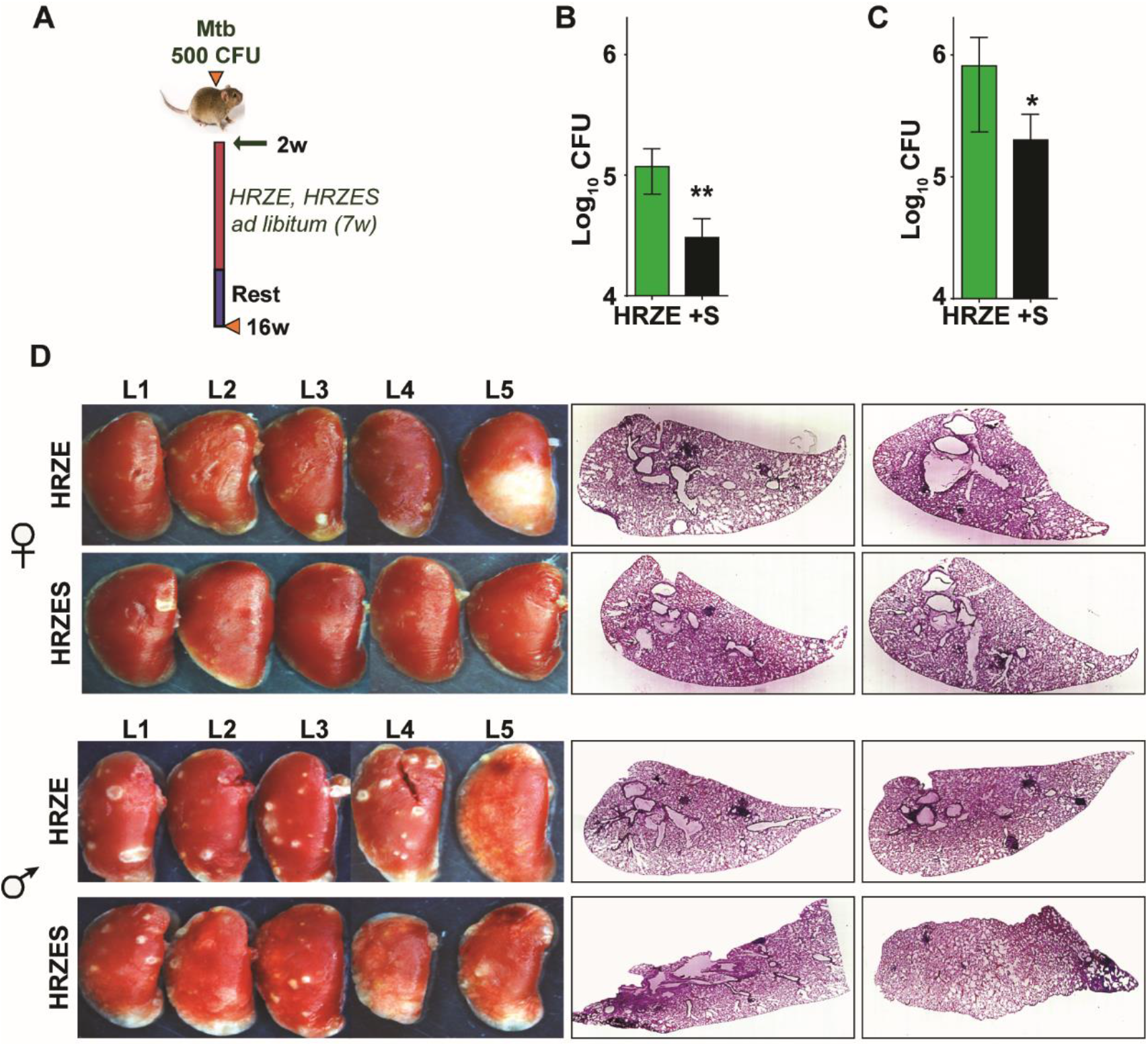
*In vivo* potentiation of SRT mediated antimycobacterial activity. A) Schematic of infected C3HeB/FeJ mice and treated with HRZE (HRC1- 1x: H-100μg/ml, R-40μg/ml, Z-150μg/ml, E-100μg/ml or in combination with SRT (10μg/ml). Animals were euthanized at the end of 16 weeks and extent of infection was determined by estimating bacterial numbers (CFU) in lungs of female (B) and male (C) mice. D) Gross lung morphology and histochemical sections of lungs with H&E staining of C3HeB/FeJ mice infected with Mtb and treated either with HRZE or in combination with SRT. (unpaired t-test,*p<0.05, **p<0.01).

To evaluate the adjunct regimen for early bacterial clearance rates, we infected C57BL/6 mice with 500 CFU of Mtb and enumerated bacterial burdens at 1- and 3-weeks post treatment according to the schedule shown in Fig. 6A. Bacterial numbers in the lungs reached ~10^7^ CFU by 4 weeks of infection (day 0 of treatment) and remained steady over the 6-week period in untreated animals (Fig. 6B). While treatment with HRZE was efficient in steadily reducing these numbers by ~100 folds, addition of SRT to the regimen significantly enhanced control by a further 2-3 folds. Moreover, the adjunct regimen was efficient in controlling dissemination of infection into spleens of infected mice (Fig. 6C). Although HRZE reduced splenic bacterial numbers significantly by 6-7 folds, HRZES was more potent reducing bacterial numbers further by ~20 folds (60-70% vs ~2-5%) by 21 days of treatment. A similar degree of enhanced bacterial control (4-6 folds lesser bacteria) was observed in lungs and spleens of Balb/c mice treated with SRT as an adjunct to conventional 4 drug-therapy (Fig. 6D). Untreated animal lungs showed a gradual consolidation of the tissue with increasing amounts of granulomatous cellular infiltration by 6 weeks of infection. Treatment with HRZE was efficient in reducing this infiltration significantly by the 6^th^ week of infection with nearly 1/6^th^ of the tissue showing signs of cellular infiltration (Fig. 6E). Lungs of mice receiving the combination showed better resolution of granulomas with significantly smaller regions of cellular collection by 3 weeks dispersed across the tissue, that was more or less absent from the tissues of mice by the 6^th^ week of treatment.

**Fig. 6:**
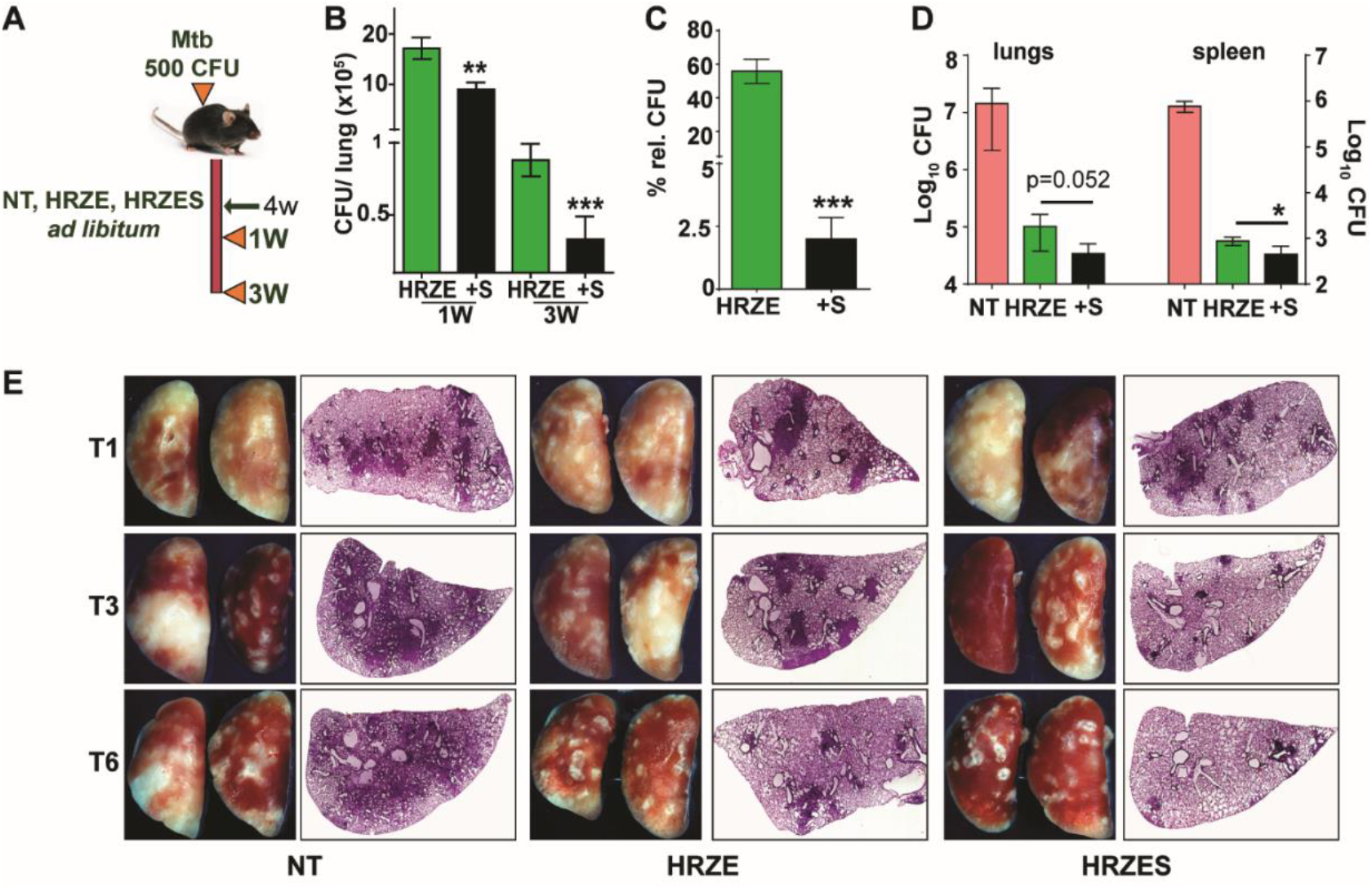
SRT initiates early bacterial clearance and dissemination in mice. A) Schematic of Mtb infection and drug treatment in C57BL6/ BalbC mice infected with Mtb and the treated with HRZE or HRZES (HRC1-1X: H-100μg/ml, R-40μg/ml, Z-150μg/ml, E-100μg/ml, SRT 10μg/ml) treatment. Lung CFU post 1 and 3 weeks (B) and spleen CFU 3 weeks after (C-relative to untreated animals) in Mtb infected C57BL6 mice of antibiotic and SRT administration. D) Lung and spleen CFUs of Balb/c mice after 3 weeks of treatment, E) Gross lung morphology and H&E staining after treatment for the indicated number of weeks. (unpaired t-test,*p<0.05, ***p<0.001).

We then tested the efficacy of the adjunct therapy against a drug tolerant Mtb strain in cellular and murine models of infection. Previously, we had demonstrated that N73- a clinical Mtb strain belonging to the L1 ancient lineage showed increasing tolerance to INH and Rifampicin as opposed to the modern L3 and L4 lineage strains by virtue of expressing the complete MmpL6 operon (60). In THP1 cells infected with Mtb, HR at C2 concentration supported stasis and failed to decrease of bacterial numbers (Fig. 7A). The combination of HR with SRT, significantly controlled infection and reduced bacterial numbers by ~10-15 folds by 3 days and ~100 folds by day5. The pattern of significantly greater bacterial control was also observed in primary human macrophages; again, a combination of HR and SRT reduced intracellular bacterial numbers by 5-50 folds in comparison to the drugs alone (Fig. 7B). With a strong indication of SRT’s ability to boost the efficacy of frontline TB drugs against tolerant Mtb, we tested its *in vivo* activity in the acute model of C3HeB/ FeJ mouse infection in combination with HR (Fig. 7C). As expected, the drugs were not efficient in controlling infection induced lesions in the lungs reducing bacterial numbers by 5 folds (Fig. 7D). However, lungs of HRS treated animals harbored ~5-8 folds lower bacterial numbers than HR treated animals. The effect of the combination was again better in controlling bacterial numbers in spleens of treated mice wherein HR did not reduce CFU in contrast to the 6-8-fold lower bacterial numbers with the combination.

**Fig. 7:**
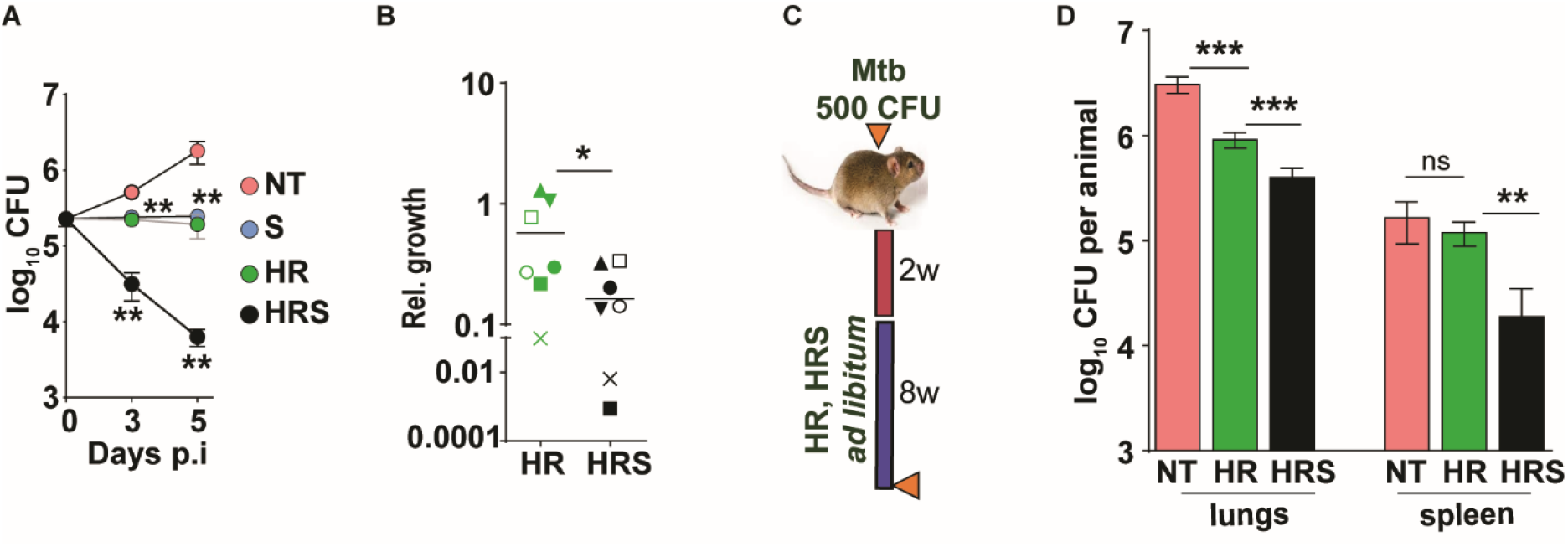
Addition of SRT helps better control of drug tolerant Mtb in vivo. A) Intracellular bacterial growth in THP1 macrophages infected with HR tolerant Mtb strain at a MOI of 5 for 6h. Following this, cells were left untreated (NT) or treated with SRT, HR or HRS for 3 or 5 days. Bacterial numbers were enumerated and is represented as average log10 CFU ± SEM from two independent experiments with triplicate wells each. B) Bacterial numbers at day 3 of primary monocyte derived macrophages (M1) from 7 independent donors. After 6h of infection, the macrophages were treated with HR and HRS for 3 days. The ratio of intracellular bacterial numbers in HR or HRS groups with respect to untreated samples is represented as relative growth with median values indicated by the horizontal line. C) Schematic of Mtb infection and drug treatment in C3HeB/FeJ mice infected with Mtb for 2 weeks followed by treatment with 0.1X HR alone or with SRT (HRS) or 8 weeks. Bacterial numbers (CFU) in lungs and spleen (D) at the end of the experiment

## DISCUSSION

Despite consistent efforts in identifying novel pathogen targeted interventions and streamlined pharmaceutical drug development control processes, fewer drugs have been accepted for clinical use in TB over the last 40 years (61). Repositioning existing drugs with established safety in humans is one of the quickest modes of developing effective control of infections that reduce the timeframe of regimen development. The need for an effective, short and pathogen-sterilizing regimen to tackle the growing problem of Mtb drug resistance and dormant bacterial populations has intensified efforts towards the development of host targeted therapies (62–65).

We and several other groups have identified type I IFN as an early response of host macrophages to infection with Mtb strains (41, 42, 49, 66). With recent evidences implicating type I IFN as a pathogen beneficial response, we hypothesized that attenuating this axis would prove beneficial in controlling bacteria in macrophages. In line with this idea, we observed that the previously reported TLR3 antagonist – sertraline (SRT) could effectively stunt Mtb induced type I IFN response in macrophages and also inhibit bacterial growth in macrophages.

SRT, along with other weak bases like fluoxetine, has been reported to moderately restrict intracellular Mtb replication in macrophages due in part to its weak basic nature without affecting the host cell viability (67). An important role for the acidic environment of bacteria resident vesicles (pH dependence) in the mycobactericidal properties of these drugs was demonstrated in this study. We also observed similar effects of SRT alone in our study- a moderate level of bacterial control in macrophages with minimal host cell death by treatment of SRT to infected macrophages. Additionally, several studies have indicated a direct action of SRT and other selective serotonin reuptake inhibitors on host immune response pathways: from enhancing the anti-inflammatory response (68), augmenting NK and CD8 cell response (69) and to inhibition of acid-sphingomyelinase (70), an essential component of the viral trafficking into NPC1+ endosomes in cells. Activation of host cell inflammasome and its antagonistic effect on the type I IFN response of cells has now been realized as important factors for controlling bacterial infections (71, 72). A direct effect on activating eicosanoids which control type I IFN following infection was identified as a key bacterial clearance mechanism of infected cells (73, 74). Consistent with this observation, targeted therapy towards elevating PGE2 activity protected mice from acute infection induced fatality (74). In line with these observations, we also describe the ability of SRT to potentiate antibiotic mediated killing by altering the inflammasome-type I IFN axis. Our results also highlight this cross-regulation between two important innate response pathways with SRT boosting the macrophage pathogen control program by repressing a pro-pathogenic and activating the host beneficial response. In line with this observation, treatment with the established inflammasome inhibitor Isoliquiritigenin reversed the ability of SRT to enhance the effects of HR in macrophages. While our work points towards the type I IFN antagonism mediated inflammasome activation as a key mechanism in providing synergy to anti TB therapy, identifying the exact molecular target of SRT in this process remains an important future challenge.

SRT combination with front-line anti-TB drugs provided early bacterial control with improved resolution of pathology and enhanced host survival, thereby being a strong candidate for host directed therapy. SRT provides additional benefits as an adjunct modality. The pharmacological properties of SRT with excellent PK-PD, safety and tolerance for long term usage in the human population has been well established (75–77). Interestingly, two patients undergoing TB therapy with INH given SRT as an anti-depressant, did not show any deleterious effects on long term use of the combination, auguring well for safety in the human population (78, 79). In addition, these studies combined with the enhanced protective capabilities of the combination therapy in pre-clinical animal models (our data), indirectly rule out any possibility of negative drug-drug interactions between SRT and ATT on prolonged usage.

The long-term standard TB treatment is associated with severe drug induced depression in patients. Recently depression was identified as an invisible co-morbidity with TB with extensive synergistic action on the patient (80, 81). It is logical to expect that SRT with its wide use as an anti-depressant in adults and children may be beneficial in tackling this dual problem efficiently with a combination regimen of frontline TB drugs and SRT. While we speculate the dual use of this adjunct treatment in treating two human disorders, detailed studies on the effective dose and duration of therapy needs to be undertaken. However, the collective properties of a SRT adjunct TB therapy - faster bacterial control, enhanced host survival and capacity to target drug tolerant/ dormant bacterial populations augurs well for the highly constrained national/ global economy combating the TB pandemic.

## Material and Methods

### Bacterial Strains and Growth Conditions

Mtb strains were cultured in Middlebrook 7H9 broth with 4% ADS or in 7H10/ 7H11 agar (BD Biosciences, USA) with 10% OADC (HiMedia laboratories, India).

### Reagents

THP1 Dual Monocytes was obtained from InvivoGen (Toulouse, France). HiglutaXL RPMI-1640 and 10% Fetal Bovine Serum (HIMedia laboratories, Mumbai, India), PMA (Phorbol 12-Mysristate 13-acetate-P8139, Sigma Aldrich, USA), BX795 (tlrl-bx7, Invivogen) were used for culture of cells. The following reagents were procured from Sigma Aldrich, USA: Vit C (L-ascorbic acid, A5960), oleic acid albumin (O3008), Isoniazid (I3377), Pyrazinamide carboxamide (P7136), Ethambutol dihydrochloride (E4630) and Sertraline hydrochloride (S6319). Rifampicin (CMS1889, HIMEDIA laboratories, Mumbai, India) and commercially available SRT (Daxid, Pfizer Ltd, India) was used for mouse studies.

### Macrophage infection

THP1 Dual reporter monocytes were grown in HiglutaXL RPMI-1640 containing 10% FBS and differentiated to macrophages with 100nM PMA for 24h. Following a period of rest for 48h, cells were infected with single cells suspensions (SCS) of Mtb at a MOI of 5 for 6h. For analyzing the Interferon (IRF pathway) activation levels, supernatants from Mtb infected THP1 Dual macrophages were assayed for stimulation by measuring luminescence as per manufacturer’s recommendations.

### Monocyte derived macrophage culture

PBMCs were isolated from fresh blood obtained from healthy donors in accordance with Institutional human ethics committee approval (Ref no: CSIR-IGIB/IHEC/2017-18 Dt. 08.02.2018). Briefly, 15-20 ml blood was collected in EDTA containing tubes and layered onto HiSep (HIMedia laboratories, Mumbai, India) and used for isolation of PBMCs according to the recommended protocols. Post RBC lysis, cells were seeded at a density of 3×10^5^cells/ well and differentiated into monocyte derived macrophages with 50ng/ml GMCSF for 7 days and then used for infection with Mtb.

### Analysis of response parameters

For analysis of different parameters of cellular response to infection, qRTPCR based gene expression analysis and cytokine ELISA in culture supernatants were performed according to manufacturer’s recommendations.

### Analysis of gene expression by qRTPCR

Total RNA was isolated from macrophages suspended in Trizol by using the recommended protocol. cDNA was prepared from 1 μg of RNA by using the Verso cDNA synthesis kit and was used at a concentration of 10ng for expression analysis by using the DyNAmo Flash SYBR Green qPCR Kit (Thermo Fisher Scientific Inc., USA).

### Analysis of cytokine secretion by ELISA

Culture supernatants at different time intervals post infection/ treatment were filtered through a 0.2μ filter and subjected to ELISA by using the eBioscience (Thermo Fisher Scientific Inc. USA) ELISA kit as per recommended protocols.

### Bacterial survival in macrophages

For determining intra cellular survival of Mtb strains macrophages were seeded in 48well plates and infected with Mtb at MOI 5 for 6 hours. SRT (20μM), BX795 (10μM) and TB drugs (C1, C2, C3: (C1-INH-200ng/ml, Rifampicin-1000ng/ml, - C2 and C3 :10- and 25-fold dilutions of C1) and used for treatment of macrophages at the appropriate concentrations. At specific days post infection macrophages were lysed with water containing 0.05% of tween80. Dilutions of the intracellular bacterial numbers were made in PBS with 0.05% of tween80 and plated on 7H10 agar plates. The VitC induced dormancy model of macrophages was developed as described earlier with cells treated with 2mM Vit C for 24h and then with Isoniazid and rifampicin for a further 3 days (53). For testing in lipid rich macrophages, cells were treated with oleic acid at 200μM concentration after PMA differentiation for 2 days, infected with Mtb for 6h, followed by treatment for 5 days and enumeration of bacterial numbers.

### Mouse infection and antibiotic treatment

(6-10 weeks old) C3HeB/FeJ/ C57BL6/ Balbc animals were infected with Mtb clinical isolate at 500 CFU per animal through aerosol route. Two/four weeks post infection animals were started on antibiotics H (100mg/l), R (40mg/l) (82), Z (150mg/l), E (100mg/l) (83) and SRT (10mg/l, human equivalent dose of 3.3 mg/kg/day), as required treatment by giving all of the drugs *ad libitum* in their drinking water for 7 weeks which was changed twice every week. For survival, animals were monitored regularly and euthanized at a pre-determined end point according to the Institutional animal ethics approval. For estimating tissue bacterial burdens, lungs and spleen of infected animals were collected in sterile saline, subjected to homogenization and used for serial dilution CFU plating on 7H11 agar plates containing OADC as supplement. Colonies were counted after incubation of the plates at 37°C for 3-5 weeks and recorded as CFU/tissue.

All statistical analysis was performed by using student’s t-test for significance, P values of < 0.05 was considered significant.

## Abbreviations

IFN: Interferon
SRT: sertraline hydrochloride
Mtb: *Mycobacterium tuberculosis*
HDT: host directed therapy

## Acknowledgements

The authors thank CSIR (VR-BSC0123), and CSIR-OLP1136 funding agency for supporting the study. CSIR-STS0016 is acknowledged for continuous maintenance of BSL3 and ABSL2 facilities. The student fellowships from CSIR, UGC and DBT India are acknowledged: DS-DBT JRF, AS-CSIR JRF, PA-CSIR-BSC0124, CSIR-SRF, SD-.DBT-JRF. The authors wish to thank Dr. Anurag Agrawal, CSIR-IGIB, New Delhi, India and Dr. Michael Glickman, MSKCC, New York, USA, for suggestions in improving the manuscript.

## Author Contributions

VR, SG, DS, PA were involved in conceptualizing and design of the work, the work was performed by DS, PA, AS and SD. The manuscript was written by VR, DS and SG with inputs from all authors.

## Conflict of interest

The authors do not have any competing interests.

## Supplementary Figure

**Fig. S1:**
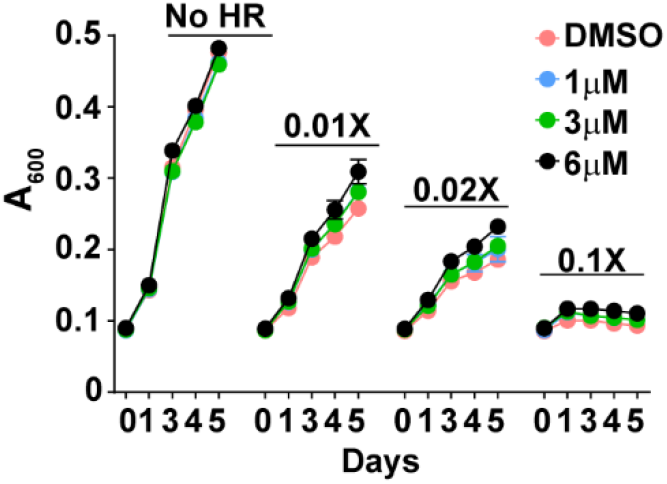
SRT does not augment the antibiotic efficacy of antibiotics *in vitro*. In vitro growth of Mtb in 7H9 media containing different concentrations of SRT. Mtb cultures were either left untreated or in the presence of different SRT and HR concentrations for 5 days at 37oC. The growth was regularly monitored by measuring the optical density of culture and is represented as mean + SD of triplicate assay wells of a representative experiment

## Notes

### Competing Interest Statement

The authors have declared no competing interest.

### Summary of Updates

WE have modified the text and included modified figures for better reading and understanding.

## References

1. Blumberg HM, et al. (2003) American Thoracic Society/Centers for Disease Control and Prevention/Infectious Diseases Society of America: treatment of tuberculosis. (Translated from eng) Am J Respir Crit Care Med 167(4):603–662 (in eng).

2. Sharma S, et al. (2017) Index-TB guidelines: Guidelines on extrapulmonary tuberculosis for India. Indian Journal of Medical Research 145(4):448–463.

3. Falzon D, et al. (2017) World Health Organization treatment guidelines for drug-resistant tuberculosis, 2016 update. (Translated from eng) Eur Respir J 49(3) (in eng).

4. Singh R, et al. (2019) Recent updates on drug resistance in Mycobacterium tuberculosis. (Translated from eng) J Appl Microbiol (in eng).

5. Hancock RE, Nijnik A, & Philpott DJ (2012) Modulating immunity as a therapy for bacterial infections. (Translated from eng) Nat Rev Microbiol 10(4):243–254 (in eng).

6. Munguia J & Nizet V (2017) Pharmacological Targeting of the Host-Pathogen Interaction: Alternatives to Classical Antibiotics to Combat Drug-Resistant Superbugs. (Translated from eng) Trends Pharmacol Sci 38(5):473–488 (in eng).

7. Parida SK, et al. (2015) T-Cell Therapy: Options for Infectious Diseases. (Translated from eng) Clin Infect Dis 61Suppl 3:S217–224 (in eng).

8. Li L, et al. (2019) Antibody Treatment against Angiopoietin-Like 4 Reduces Pulmonary Edema and Injury in Secondary Pneumococcal Pneumonia. (Translated from eng) MBio 10(3) (in eng).

9. Foster GR (2010) Pegylated interferons for the treatment of chronic hepatitis C: pharmacological and clinical differences between peginterferon-alpha-2a and peginterferon-alpha-2b. (Translated from eng) Drugs 70(2):147–165 (in eng).

10. Zakaria MK, Carletti T, & Marcello A (2018) Cellular Targets for the Treatment of Flavivirus Infections. (Translated from eng) Front Cell Infect Microbiol 8:398 (in eng).

11. Fedson DS, Jacobson JR, Rordam OM, & Opal SM (2015) Treating the Host Response to Ebola Virus Disease with Generic Statins and Angiotensin Receptor Blockers. (Translated from eng) MBio 6(3):e00716 (in eng).

12. Skerry C, Scanlon K, Rosen H, & Carbonetti NH (2015) Sphingosine-1-phosphate Receptor Agonism Reduces Bordetella pertussis-mediated Lung Pathology. (Translated from eng) J Infect Dis 211(12):1883–1886 (in eng).

13. Scanlon KM, Skerry C, & Carbonetti NH (2015) Novel therapies for the treatment of pertussis disease. (Translated from eng) Pathog Dis 73(8):ftv074 (in eng).

14. Yedery RD & Jerse AE (2015) Augmentation of Cationic Antimicrobial Peptide Production with Histone Deacetylase Inhibitors as a Novel Epigenetic Therapy for Bacterial Infections. (Translated from eng) Antibiotics (Basel) 4(1):44–61 (in eng).

15. Jimenez de Oya N, Blazquez AB, Casas J, Saiz JC, & Martin-Acebes MA (2018) Direct Activation of Adenosine Monophosphate-Activated Protein Kinase (AMPK) by PF-06409577 Inhibits Flavivirus Infection through Modification of Host Cell Lipid Metabolism. (Translated from eng) Antimicrob Agents Chemother 62(7) (in eng).

16. Lange SM, et al. (2019) l-Arginine Synthesis from l-Citrulline in Myeloid Cells Drives Host Defense against Mycobacteria In Vivo. (Translated from eng) J Immunol 202(6):1747–1754 (in eng).

17. Phelan JJ, et al. (2018) Modulating Iron for Metabolic Support of TB Host Defense. (Translated from eng) Front Immunol 9:2296 (in eng).

18. Oh KB, Oh MN, Kim JG, Shin DS, & Shin J (2006) Inhibition of sortase-mediated Staphylococcus aureus adhesion to fibronectin via fibronectin-binding protein by sortase inhibitors. (Translated from eng) Appl Microbiol Biotechnol 70(1):102–106 (in eng).

19. Cusumano CK, et al. (2011) Treatment and prevention of urinary tract infection with orally active FimH inhibitors. (Translated from eng) Sci Transl Med 3(109):109ra115 (in eng).

20. Berube BJ & Bubeck Wardenburg J (2013) Staphylococcus aureus alpha-toxin: nearly a century of intrigue. (Translated from eng) Toxins (Basel) 5(6):1140–1166 (in eng).

21. Blum CA, et al. (2015) Adjunct prednisone therapy for patients with community-acquired pneumonia: a multicentre, double-blind, randomised, placebo-controlled trial. (Translated from eng) Lancet 385(9977):1511–1518 (in eng).

22. Benmerzoug S, et al. (2018) GM-CSF targeted immunomodulation affects host response to M. tuberculosis infection. (Translated from eng) Sci Rep 8(1):8652 (in eng).

23. de Martino M, Lodi L, Galli L, & Chiappini E (2019) Immune Response to Mycobacterium tuberculosis: A Narrative Review. (Translated from eng) Front Pediatr 7:350 (in eng).

24. Shi L, Jiang Q, Bushkin Y, Subbian S, & Tyagi S (2019) Biphasic Dynamics of Macrophage Immunometabolism during Mycobacterium tuberculosis Infection. (Translated from eng) MBio 10(2) (in eng).

25. Liu CH, Liu H, & Ge B (2017) Innate immunity in tuberculosis: host defense vs pathogen evasion. (Translated from eng) Cell Mol Immunol 14(12):963–975 (in eng).

26. Padhi A, et al. (2019) Mycobacterium tuberculosis LprE Suppresses TLR2-Dependent Cathelicidin and Autophagy Expression to Enhance Bacterial Survival in Macrophages. (Translated from eng) J Immunol 203(10):2665–2678 (in eng).

27. Dey RJ, et al. (2017) Inhibition of innate immune cytosolic surveillance by an M. tuberculosis phosphodiesterase. (Translated from eng) Nat Chem Biol 13(2):210–217 (in eng).

28. Gupta S, Winglee K, Gallo R, & Bishai WR (2017) Bacterial subversion of cAMP signalling inhibits cathelicidin expression, which is required for innate resistance to Mycobacterium tuberculosis. (Translated from eng) J Pathol 242(1):52–61 (in eng).

29. Brites D & Gagneux S (2015) Co-evolution of Mycobacterium tuberculosis and Homo sapiens. (Translated from eng) Immunol Rev 264(1):6–24 (in eng).

30. Lin W, et al. (2016) Transcriptional Profiling of Mycobacterium tuberculosis Exposed to In Vitro Lysosomal Stress. (Translated from eng) Infect Immun 84(9):2505–2523 (in eng).

31. Smith DG, et al. (2019) Identification and characterization of a novel anti-inflammatory lipid isolated from Mycobacterium vaccae, a soil-derived bacterium with immunoregulatory and stress resilience properties. (Translated from eng) Psychopharmacology (Berl) 236(5):1653–1670 (in eng).

32. Mehta M & Singh A (2019) Mycobacterium tuberculosis WhiB3 maintains redox homeostasis and survival in response to reactive oxygen and nitrogen species. (Translated from eng) Free Radic Biol Med 131:50–58 (in eng).

33. Salahuddin N, et al. (2013) Vitamin D accelerates clinical recovery from tuberculosis: results of the SUCCINCT Study [Supplementary Cholecalciferol in recovery from tuberculosis]. A randomized, placebo-controlled, clinical trial of vitamin D supplementation in patients with pulmonary tuberculosis’. (Translated from eng) BMC Infect Dis 13:22 (in eng).

34. Suarez-Mendez R, et al. (2004) Adjuvant interferon gamma in patients with drug - resistant pulmonary tuberculosis: a pilot study. (Translated from eng) BMC Infect Dis 4:44 (in eng).

35. Singhal A, et al. (2014) Metformin as adjunct antituberculosis therapy. (Translated from eng) Sci Transl Med 6(263):263ra159 (in eng).

36. Naftalin CM, et al. (2018) Adjunctive use of celecoxib with anti-tuberculosis drugs: evaluation in a whole-blood bactericidal activity model. (Translated from eng) Sci Rep 8(1):13491 (in eng).

37. Lachmandas E, et al. (2019) Metformin Alters Human Host Responses to Mycobacterium tuberculosis in Healthy Subjects. (Translated from eng) J Infect Dis 220(1):139–150 (in eng).

38. Rayasam GV & Balganesh TS (2015) Exploring the potential of adjunct therapy in tuberculosis. (Translated from eng) Trends Pharmacol Sci 36(8):506–513 (in eng).

39. Dorhoi A, et al. (2014) Type I IFN signaling triggers immunopathology in tuberculosis-susceptible mice by modulating lung phagocyte dynamics. (Translated from eng) Eur J Immunol 44(8):2380–2393 (in eng).

40. Watson RO, et al. (2015) The Cytosolic Sensor cGAS Detects Mycobacterium tuberculosis DNA to Induce Type I Interferons and Activate Autophagy. (Translated from eng) Cell Host Microbe 17(6):811–819 (in eng).

41. Donovan ML, Schultz TE, Duke TJ, & Blumenthal A (2017) Type I Interferons in the Pathogenesis of Tuberculosis: Molecular Drivers and Immunological Consequences. (Translated from eng) Front Immunol 8:1633 (in eng).

42. Shankaran D, Arumugam P, Bothra A, Gandotra S, & Vivek R (bioRxiv.

43. Zhu J, et al. (2010) High-throughput screening for TLR3-IFN regulatory factor 3 signaling pathway modulators identifies several antipsychotic drugs as TLR inhibitors. (Translated from eng) J Immunol 184(10):5768–5776 (in eng).

44. Liu CH, Liu H, & Ge B (2017) Innate immunity in tuberculosis: host defense vs pathogen evasion. (Translated from eng) Cell Mol Immunol (in eng).

45. Xu G, Wang J, Gao GF, & Liu CH (2014) Insights into battles between Mycobacterium tuberculosis and macrophages. (Translated from eng) Protein Cell 5(10):728–736 (in eng).

46. Petit-Jentreau L, Tailleux L, & Coombes JL (2017) Purinergic Signaling: A Common Path in the Macrophage Response against Mycobacterium tuberculosis and Toxoplasma gondii. (Translated from eng) Front Cell Infect Microbiol 7:347 (in eng).

47. Wassermann R, et al. (2015) Mycobacterium tuberculosis Differentially Activates cGAS- and Inflammasome-Dependent Intracellular Immune Responses through ESX-1. (Translated from eng) Cell Host Microbe 17(6):799–810 (in eng).

48. Wiens KE & Ernst JD (2016) The Mechanism for Type I Interferon Induction by Mycobacterium tuberculosis is Bacterial Strain-Dependent. (Translated from eng) PLoS Pathog 12(8):e1005809 (in eng).

49. Moreira-Teixeira L, Mayer-Barber K, Sher A, & O’Garra A (2018) Type I interferons in tuberculosis: Foe and occasionally friend. (Translated from eng) J Exp Med 215(5):1273–1285 (in eng).

50. Cicchese JM, Dartois V, Kirschner DE, & Linderman JJ (2020) Both Pharmacokinetic Variability and Granuloma Heterogeneity Impact the Ability of the First-Line Antibiotics to Sterilize Tuberculosis Granulomas. (Translated from eng) Front Pharmacol 11:333 (in eng).

51. Prideaux B, et al. (2015) The association between sterilizing activity and drug distribution into tuberculosis lesions. (Translated from eng) Nat Med 21(10):1223–1227 (in eng).

52. Sarathy JP, et al. (2018) Extreme Drug Tolerance of Mycobacterium tuberculosis in Caseum. (Translated from eng) Antimicrob Agents Chemother 62(2) (in eng).

53. Sikri K, et al. (2018) Multifaceted remodeling by vitamin C boosts sensitivity of Mycobacterium tuberculosis subpopulations to combination treatment by anti-tubercular drugs. (Translated from eng) Redox Biol 15:452–466 (in eng).

54. Dartois V (2014) The path of anti-tuberculosis drugs: from blood to lesions to mycobacterial cells. (Translated from eng) Nat Rev Microbiol 12(3):159–167 (in eng).

55. Jaisinghani N, et al. (2018) Necrosis Driven Triglyceride Synthesis Primes Macrophages for Inflammation During Mycobacterium tuberculosis Infection. (Translated from eng) Front Immunol 9:1490 (in eng).

56. Teles RM, et al. (2013) Type I interferon suppresses type II interferon-triggered human anti-mycobacterial responses. (Translated from eng) Science 339(6126):1448–1453 (in eng).

57. Sitges M, Gomez CD, & Aldana BI (2014) Sertraline reduces IL-1beta and TNF-alpha mRNA expression and overcomes their rise induced by seizures in the rat hippocampus. (Translated from eng) PLoS One 9(11):e111665 (in eng).

58. Szalach LP, Lisowska KA, & Cubala WJ (2019) The Influence of Antidepressants on the Immune System. (Translated from eng) Arch Immunol Ther Exp (Warsz) 67(3):143–151 (in eng).

59. Ruiz-Grosso P, Cachay R, de la Flor A, Schwalb A, & Ugarte-Gil C (2020) Association between tuberculosis and depression on negative outcomes of tuberculosis treatment: A systematic review and meta-analysis. (Translated from eng) PLoS One 15(1):e0227472 (in eng).

60. Arumugam P, Shankaran D, Bothra A, Gandotra S, & Rao V (2019) The MmpS6-MmpL6 Operon Is an Oxidative Stress Response System Providing Selective Advantage to Mycobacterium tuberculosis in Stress. (Translated from eng) J Infect Dis 219(3):459–469 (in eng).

61. Goel D (2014) Bedaquiline: A novel drug to combat multiple drug-resistant tuberculosis. (Translated from eng) J Pharmacol Pharmacother 5(1):76–78 (in eng).

62. Palucci I & Delogu G (2018) Host Directed Therapies for Tuberculosis: Futures Strategies for an Ancient Disease. (Translated from eng) Chemotherapy 63(3):172–180 (in eng).

63. Mishra R, et al. (2019) Targeting redox heterogeneity to counteract drug tolerance in replicating Mycobacterium tuberculosis. (Translated from eng) Sci Transl Med 11(518) (in eng).

64. Fatima S, et al. (2019) Mycobacterium tuberculosis programs mesenchymal stem cells to establish dormancy and persistence. (Translated from eng) J Clin Invest (in eng).

65. Dara Y, Volcani D, Shah K, Shin K, & Venketaraman V (2019) Potentials of Host-Directed Therapies in Tuberculosis Management. (Translated from eng) J Clin Med 8(8) (in eng).

66. McNab FW, et al. (2014) Type I IFN induces IL-10 production in an IL-27-independent manner and blocks responsiveness to IFN-gamma for production of IL-12 and bacterial killing in Mycobacterium tuberculosis-infected macrophages. (Translated from eng) J Immunol 193(7):3600–3612 (in eng).

67. Schump MD, Fox DM, Bertozzi CR, & Riley LW (2017) Subcellular Partitioning and Intramacrophage Selectivity of Antimicrobial Compounds against Mycobacterium tuberculosis. (Translated from eng) Antimicrob Agents Chemother 61(3) (in eng).

68. Hannestad J, DellaGioia N, & Bloch M (2011) The effect of antidepressant medication treatment on serum levels of inflammatory cytokines: a meta-analysis. (Translated from eng) Neuropsychopharmacology 36(12):2452–2459 (in eng).

69. Benton T, et al. (2010) Selective serotonin reuptake inhibitor suppression of HIV infectivity and replication. (Translated from eng) Psychosom Med 72(9):925–932 (in eng).

70. Kouznetsova J, et al. (2014) Identification of 53 compounds that block Ebola virus-like particle entry via a repurposing screen of approved drugs. (Translated from eng) Emerg Microbes Infect 3(12):e84 (in eng).

71. Novikov A, et al. (2011) Mycobacterium tuberculosis triggers host type I IFN signaling to regulate IL-1beta production in human macrophages. (Translated from eng) J Immunol 187(5):2540–2547 (in eng).

72. Ji DX, et al. (2019) Type I interferon-driven susceptibility to Mycobacterium tuberculosis is mediated by IL-1Ra. (Translated from eng) Nat Microbiol 4(12):2128–2135 (in eng).

73. Hawn TR, Shah JA, & Kalman D (2015) New tricks for old dogs: countering antibiotic resistance in tuberculosis with host-directed therapeutics. (Translated from eng) Immunol Rev 264(1):344–362 (in eng).

74. Mayer-Barber KD, et al. (2014) Host-directed therapy of tuberculosis based on interleukin-1 and type I interferon crosstalk. (Translated from eng) Nature 511(7507):99–103 (in eng).

75. Sheehan DV & Kamijima K (2009) An evidence-based review of the clinical use of sertraline in mood and anxiety disorders. (Translated from eng) Int Clin Psychopharmacol 24(2):43–60 (in eng).

76. Ronfeld RA, Wilner KD, & Baris BA (1997) Sertraline. Chronopharmacokinetics and the effect of coadministration with food. (Translated from eng) Clin Pharmacokinet 32 Suppl 1:50–55 (in eng).

77. Mandrioli R, Mercolini L, & Raggi MA (2013) Evaluation of the pharmacokinetics, safety and clinical efficacy of sertraline used to treat social anxiety. (Translated from eng) Expert Opin Drug Metab Toxicol 9(11):1495–1505 (in eng).

78. Malek-Ahmadi P, Chavez M, & Contreras SA (1996) Coadministration of isoniazid and antidepressant drugs. (Translated from eng) J Clin Psychiatry 57(11):550 (in eng).

79. Judd FK, Mijch AM, Cockram A, & Norman TR (1994) Isoniazid and antidepressants: is there cause for concern? (Translated from eng) Int Clin Psychopharmacol 9(2):123–125 (in eng).

80. Sweetland AC, et al. (2017) Addressing the tuberculosis-depression syndemic to end the tuberculosis epidemic. (Translated from eng) Int J Tuberc Lung Dis 21(8):852–861 (in eng).

81. Trenton AJ & Currier GW (2001) Treatment of Comorbid Tuberculosis and Depression. (Translated from eng) Prim Care Companion J Clin Psychiatry 3(6):236–243 (in eng).

82. Vilcheze C, Kim J, & Jacobs WR, Jr. (2018) Vitamin C Potentiates the Killing of Mycobacterium tuberculosis by the First-Line Tuberculosis Drugs Isoniazid and Rifampin in Mice. (Translated from eng) Antimicrob Agents Chemother 62(3) (in eng).

83. Lanoix JP, Betoudji F, & Nuermberger E (2016) Sterilizing Activity of Pyrazinamide in Combination with First-Line Drugs in a C3HeB/FeJ Mouse Model of Tuberculosis. (Translated from eng) Antimicrob Agents Chemother 60(2):1091–1096 (in eng).

